# Bimodal peptide collision cross section distribution reflects two stable conformations in the gas phase

**DOI:** 10.1101/2025.05.19.654929

**Authors:** Juan Restrepo, Daniel Szoelloesi, Tobias Kiermeyer, Christoph Wichmann, Helmut Grubmüller, Jürgen Cox

## Abstract

Recent high throughput applications to shotgun proteomics have shown great benefits of coupling ion mobility spectrometry (IMS) to mass spectrometry. IMS adds a separation dimension by differentiating biomolecules by their size and shape. We (and others) find that the distribution of peptide collision cross section (CCS) is often bimodal, which limits the utility of current machine learning predictions for peptide identification. Molecular dynamics simulations indicate that the peptides in the drift tube can adopt multiple stable conformations and that the two modes correspond to predominantly extended (mostly helical) and more compact (globular and less ordered) conformations. Most peptides have a charge-dependent strong preference for one of the two conformations, while some can adapt both, as evidenced by a simple geometric model of the CCS data. We suggest a novel two-valued CCS predictor allowing for multiple peptide conformations. Its integration into data-independent acquisition proteomics increases identification rates of peptides compared to single-value predictors.

## INTRODUCTION

Ion mobility spectrometry^1–3^ (IMS) is a method for separating ionized molecules in the gas phase based on their mobility in a carrier gas. Measured ion mobility values can be converted to rotationally averaged collision cross sections (CCS)^4^, which are correlated to the three-dimensional structure of the ionized molecules. Therefore, it can separate molecules by their sizes and conformations in complex samples. Shotgun proteomics^5–8^ involves measuring more than 100,000 different peptides from a single liquid chromatography-mass spectrometry (LC-MS) run^9^. Here, using IMS after liquid chromatography has proven to be beneficial^10–13^ because the separation of co-eluting peptides by their CCS leads to less complex mass spectra and subsequent benefits in their identification^14^. Besides the reduced complexity, the MS features get annotated with their CCS values when LC-MS is coupled to IMS. This additional data dimension can be used to reduce the search space during peptide identification and thereby increase the reliability of identification confidence. However, unlike the sequences, which are known *a priori* and are available in databases for searching, the CCS values of ionized peptides in the gas phase are in general unknown, since the three-dimensional structures of gas-phase peptides are experimentally and computationally hard to obtain, particularly at large scale. Furthermore, it is known that peptides can produce complex ion mobility spectra (mobilograms) with multiple peaks. This can occur due to multiple energetically favorable configurations in the gas phase^15^ or other more complex mechanisms^16,17,18,19^. In either case, the mobilogram can serve as a rich source of physical information for identification.

The conformations of ionized peptides in the gas phase have been studied both experimentally and theoretically, the latter mainly by molecular dynamics (MD) simulations. Many studies focused on Alanine-based peptides^20–24^ and have shown that different mobility values represent specific peptide conformations such as alpha helices, hinged helix-coils, globular, and open globular structures. Furthermore, alpha helix stability has been studied in a wide range of experimental conditions^21,23,24^ as well as its unfolding dynamics^22^. These results not only show that ionized peptides can have various stable conformations, but also prove that MD simulations can offer a toolbox for understanding IM experiments on the atomic level. Nevertheless, the molecular mechanisms leading to the various peptide conformations as well as their interconversion are highly complex and remain only partially understood. Breuker & McLafferty^25^ showed that proteins retain water molecules during the electrospray ionization and keep the initial condensed phase structures in the gas phase. However, these water molecules are not detected in a large-scale proteomics experiment. Subsequent studies on the stability of these structures showed that helices tend to unfold into globular structures when the temperature of the ion is increased^16,26^ and also when they pass through a drift tube filled with a buffer gas and an electric field^22^.

To develop a physically sound model for CCS values of peptides it is therefore crucial to better understand the underlying physical reasons of the complexity in the IMS mobilograms of peptides and to incorporate these findings into the model. To this end, it is also essential to gain a more fundamental understanding of the molecular scattering processes that give rise to the observed CCS values. Current prediction approaches use either regression on sequence space, as they can be applied to retention time prediction as well^27^, or empirically driven models^14,28,29^ that explicitly parameterize the influence of amino acid positions on size and structure. These approaches usually neglect that a peptide molecule can have more than one CCS value or, more recently^30^, allow for several values without representing the actual peptide dynamics and not much impact on identifications. Furthermore, benchmarking predictions on pre-filtered data with single valued CCS, might not be a realistic setting, since the actual information available in the identification process is more complex.

Here we want to reveal the structural origin of the bimodal behavior of ion mobility values observed in peptide populations and find a model for the IMS values of ionized peptides. To this end, we combined analysis of a large-scale LC-IMS-MS/MS shotgun proteomics dataset with molecular dynamics simulations of *in vacuo* folding of representative peptides, as well as of their motion through the diluted gas within the drift tube. Our MD results suggest the two modes correspond to distinct conformations: one globular (compact) and the other helical (extended), with some peptides being able to adopt either state. Our simulations also reveal three different modes of how the peptide interacts with individual gas molecules, which, combined, determine the terminal drift velocity and hence its mobility. To extend these findings beyond computationally expensive MD simulations, we applied further computational and statistical methods. Starting with an idealized scattering model of helices and spheres for qualitative insights, we then developed a geometric model that accurately fits measured CCS values, providing a quantitative description of the bimodal distribution. Lastly, we developed a performance-driven method which focuses on separating the two populations as well as possible to enable the development of a novel, multi-valued CCS prediction method and benchmarked with respect to its impact on the number of identified peptides.

## RESULTS

### Bi-modality of peptide collision cross section distribution

We re-analyzed a large LC-IMS-MS/MS shotgun proteomics dataset^33^ spanning five species and the three proteases Trypsin, LysC and LysN. Fig. 1a-c shows the distribution of peptides in the trypsin-digested dataset in the space of reduced mobility vs. mass-to-charge ratio. Two subpopulations which overlapped with each other are observed with three positive charges over the m/z-1/K0 plane as shown in Fig. 1b. A fit to the sum of two bivariate normal distributions (Supplementary Fig. 1b) assigns 55 and 45 percent of data points to the upper and lower cloud, respectively. Separating the data by species (Supplementary Fig. 2) shows no significant inter-species differences in the distributions of data points, suggesting that the observed bimodality arises from physio-chemical effects independent of species origin. Examination of the same charge state across proteases in Fig. 1 reveals a consistent trend: as the net charge increases, a higher proportion of peptides is observed in the upper population across all enzymes. Fig. 1g presents the estimated percentages of peptides in the lower population for all combinations, further corroborating this trend.

**Figure 1:**
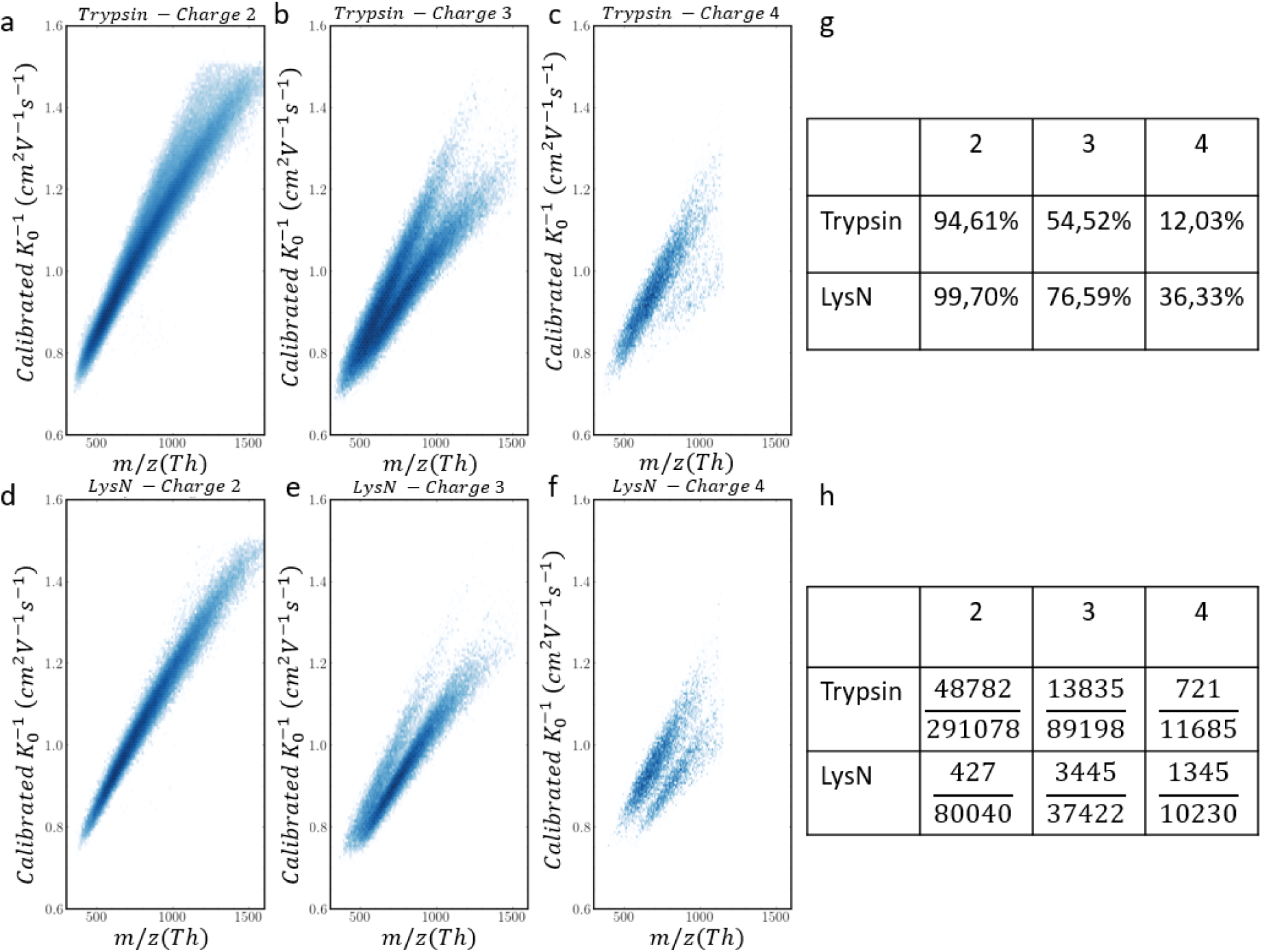
Bimodal distribution. **a.-f.** Distribution of CCS vs, m/z values across proteases Trypsin and LysN and net charges +2, +3 and +4. **g.** Estimated percentages of peptides in the lower population. **h.** Number of peptides identified in both populations for a given charge state and protease.

Additionally, analysis of the different charge states per protease indicates that, for all charge states, peptides digested with Trypsin and LysC (Supplementary Fig. 3cgk) exhibit a stronger preference for the upper population compared to those digested with LysN (Fig. 1d-f). By looking at the net charge of the termini (Supplementary Fig. 4), it is also clear that peptides digested with LysN and with positively charged amino acids close to the C-terminus tend to be in the upper population while negatively charged ones tend to be in the lower population. A similar behavior but with lower intensity can also be seen for peptides digested with LysC/Trypsin. Peptides can be found in both populations, Fig. 1h shows the ratio of such peptides, and even occur in more than two versions, which results in complex 1/K0 spectra (Supplementary Fig. 5). Previous studies have examined the link between peptide sequence and the two subpopulations, but, although significant enrichments exist, the effects are small and do not fully explain the bimodality. We therefore have approached the question why some peptides have multiple ion mobilities and how the observed CCS values arise from accumulated individual collisions with gas molecules in the drift tube from first principles using atomistic molecular dynamics (MD) simulations. In particular, to answer the question why some peptides have multiple ion mobilities, we asked (i) can the same peptide adopt multiple conformations in the drift tube environment?, and (ii) does the conformation affect the drift velocity and of so, how? To tackle these problems, we carried out MD simulations with 12 sample peptides selected from the large-scale dataset (Table 1).

**Table 1.**
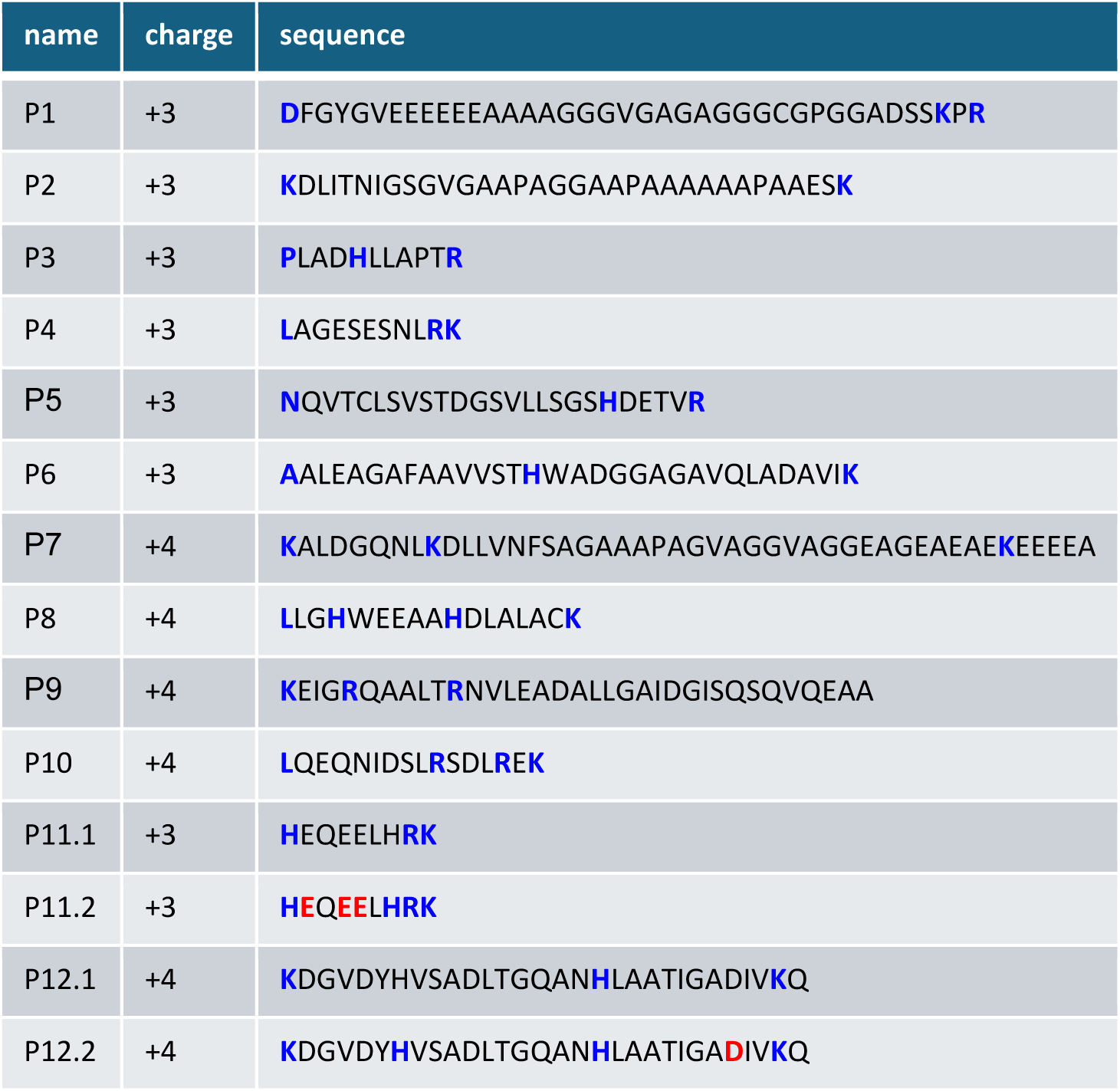
Tested peptide sequences, total charge and charge location. Positively and negatively charged residues are shown as bold blue and red.

### MD simulations reveal a strong prevalence of globular and helical peptide conformations in vacuum

Conformations *in vacuo* were determined by ‘temperature quenching’ MD simulations. All simulations were started at a temperature of 600 K, which was subsequently gradually decreased to 305 K over a period of 0.5 µs. Each of these simulations was initiated from a fully extended peptide conformation, mimicking high temperature release from the solvent and subsequent conformational quenching in the gas phase, as expected to occur in the experiments. For each of the selected 12 peptides, 1000 such quenching simulations were carried out. The changes in the peptide conformation were characterized through CCS values predicted from simulation frames by the software IMoS, as well as by their helix content.

Figure 2a shows two example structures of the P1 peptide, a largely helical one with a relatively large predicted CCS value, and a more globular structure with higher variability, no preferred fold, and with a smaller predicted CCS. Fig. 2b shows the evolution of the 1000 simulations for the P1 peptide as the temperature decreases. Initially, high temperature conformations with large CCS dominate, whereas with decreasing temperature two markedly lower CCS populations at 750 and 1000 Å^2^ emerge, the distribution of which converges towards the end of the simulations (Fig. 2c). Notably, a clear gap between the two main CCS populations is seen at about 850 Å^2^, and exchange between these ceases at already 490 K. The two main conformations shown in Fig. 2a are seen also for other peptides, e.g. P5, as shown by their final CCS distributions (Fig. 2d). For comparison, the CCS values measured in the experiment are indicated as red markers. The values predicted from MD structures follow the same trend as the measured CCS, but differ by an offset, in line with the observation by Ewing et al.^31^, according to which the predicted CCS values are about 5% larger than the measured ones.

**Figure 2:**
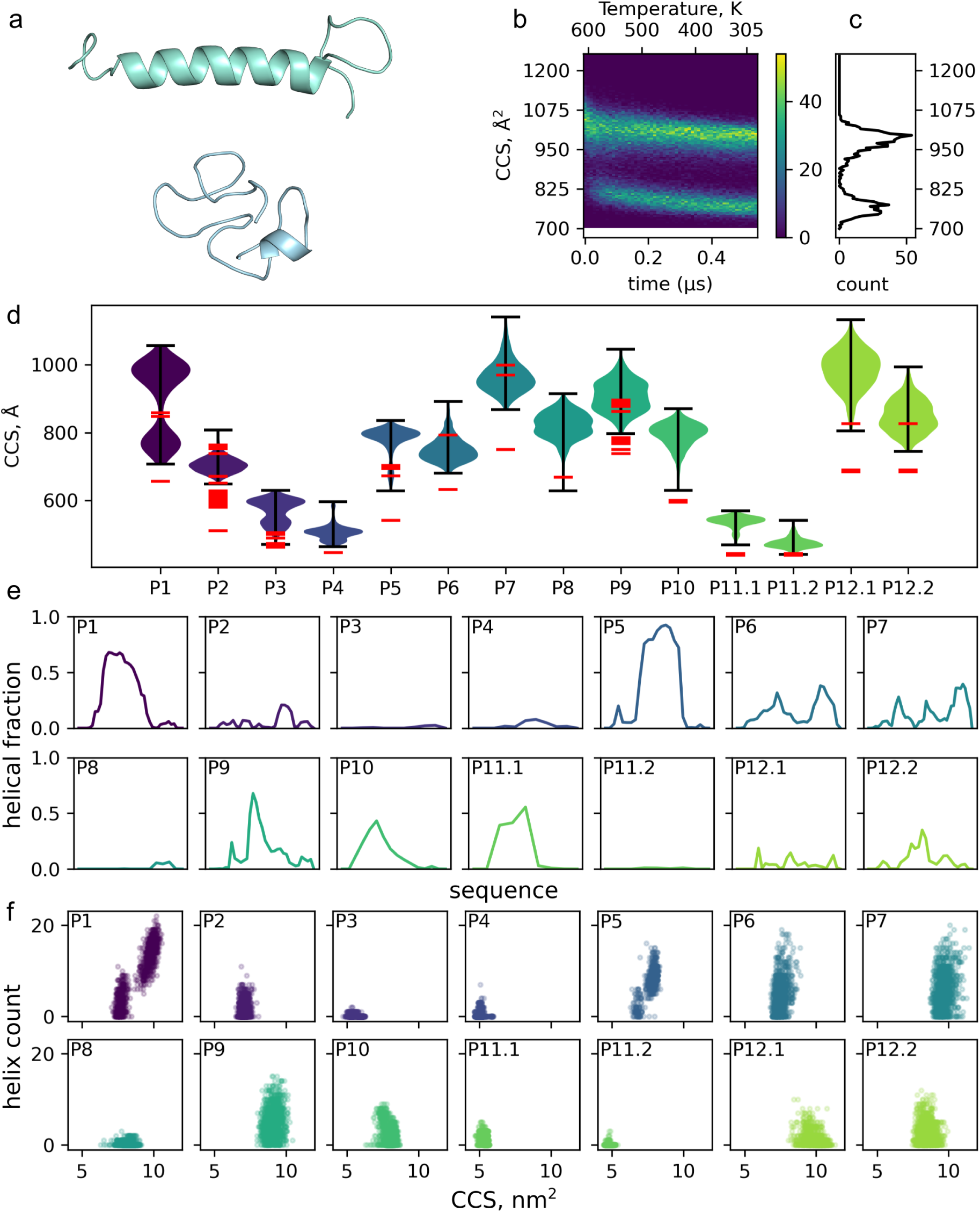
Peptide conformation in vacuum. **a.** Example conformations of the two major folds. **b.** Evolution of the CCS over the quenching simulation and **c.** CCS distribution of the final conformations. **d.** Violin plot of CCS for all simulated peptides. Peptide P11 and P12 was tested with two different charge distributions indicated by the label. Red line segments indicate available measurement data (peak positions). **e.** The fraction of helical residue along the tested sequence compared to the 1000 quenching simulations using the final conformation. **f.** Count of helical residues versus predicted CCS using the final conformation of the quenching simulations.

As can also be seen in Fig. 2d, several peptides indeed show multiple CSS values even for the same charge distribution, with P1 as the most incisive example. Of the other peptides, many also show wide CCS distributions, indicating more than one conformation. For P11 and P12, where two different charge distributions were tested (P11.1 vs. P11.2 and P12.1 vs. P12.2), the resulting distributions differ, too, albeit both charge states sample the whole CCS space. This observation suggests that, in addition to the different conformations observed for identical charge states, alternative charge states can add to the multimodality of the observed CCS distibutions.

The example of P1 indicates that α-helices can and do form *in vacuo,* which is plausible due its hydrophobicity, akin to membrane interiors, which also promote the formation of α-helices. For all 12 peptides, Figs. 2e and f summarize the abundance of α-helical content along the sequence (Fig. 2e) as well as how the predicted CCS relates to the helical content (Fig. 2f. Because some of the unrelated peptides show considerable helical content, and nearly all of them some, we conclude that helix formation is a general phenomenon, and may indeed occur for a larger fraction of all measured peptides.

Notably, although the fraction of helical residues generally correlates with the CCS for the larger peptides (Fig. 2f), the correlation is much weaker for the smaller peptides. Closer inspection of the respective structures shows that this is because the CCS values of a short helix and a globular fold are rather similar, which also explains why the two populations merge at low peptide mass.

To assess whether or not the observed conformations are kinetically trapped or, rather, in thermal equilibrium, Supplementary Fig. 6 shows the potential energy distributions of P1 in its two final conformations during last 40 ns of the quenching simulations at constant temperature 305 K. The distributions are largely overlapping, with an only small difference between the respective average potential energies between the two conformations. This indicates that the two conformations are indeed close to thermal equilibrium and, therefore, their final distributions are expected to be rather insensitive to the chosen cooling rates, the precise value of which in the experiments is unknown.

### MD simulations of peptides within the drift tube environment show that conformational differences suffice for drift velocity bimodality

Are the structural differences predicted by the MD quenching simulations large enough to explain the measured bimodal ion mobility distributions? To answer this question, we performed fully atomistic MD simulations of P1, for which the globular and helical conformations are well separated, within the drift tube environment including the electric field that accelerates the peptide (see methods) as well as the opposing air resistance due to collisions with the gas molecules. Here we did not want to resort to established structure-based CCS estimates, because it is unknown (a) to what extent the peptide conformation is changed due to the ‘bombardment’ by the gas molecules within the drift tube and (b) what the nature of the collisions is that ultimately determine the effective CCS. Our new type of MD simulations served to also address these questions.

From the quenching simulations seven different globular and seven helical conformations were selected; each of these was placed within 10 different boxes of 10^6^ nm^3^ each, filled with 2.7 mbar air (51 N_2_ and 13 O_2_ molecules) with different random gas positions and velocities, resulting in a total of 140 simulations. In each of the simulations, the peptides started at rest, and – similar to the experiment, – an electric field of 20 V/cm was applied to accelerate the peptide while the center of mass of the gas molecules was kept stationary. Due to the electric field, the peptide gradually accelerated while colliding with the gas molecules. In order to maintain the gas temperature but do not perturb peptide velocities, only the gas was coupled to a heat bath (see Online methods). Visual inspection of the simulations with high temporal resolution (1 frame/ps) revealed three main collision types sketched in Fig. 3a and visualized in the Supplementary movies 1-4. The first type, the expected one, is mostly elastic, with a very short interaction time; second, and unexpectedly, we observed adsorption with subsequent reemission, where the gas molecule spent an extended time span (up to 3.5 ns) on the surface of the peptide; third, swing-by events, during which, in contrast to a collision, the protein and gas interact attractively through Lennard-Jones forces and thus also change velocities. This third type of collision was also unexpected due to the weakness of the Lennard-Jones interactions.

**Figure 3.**
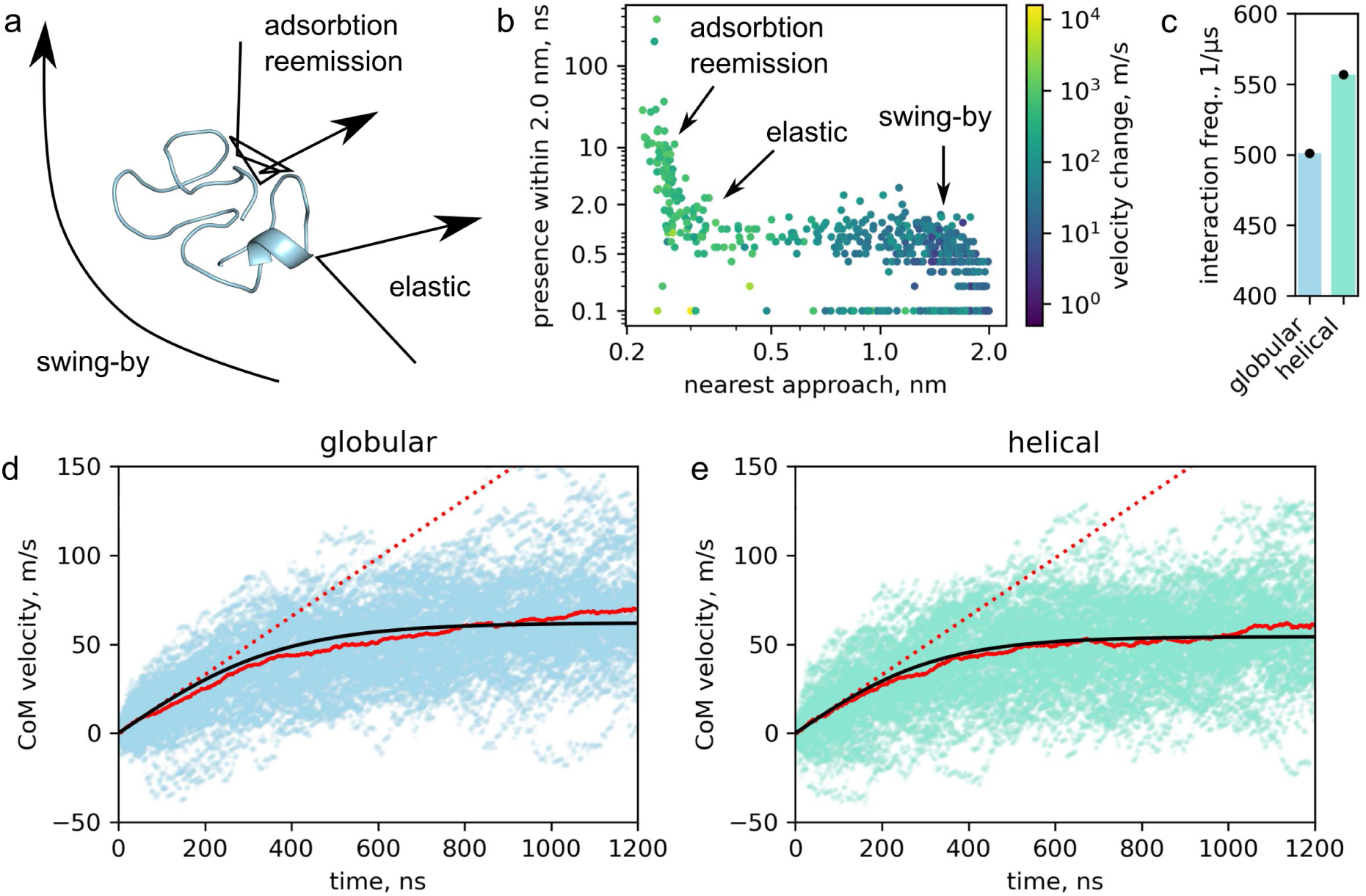
Collisions and simulated drift velocity. **a.** Scheme of the three main collision types. **b.** Collision events are tested for nearest proximity, time spent within 2 nm and the change of the velocity before and after the event. The location of the tree collision types are highlighted **c.** The number of interactions/collisions averaged for the globular and helical folds in the drift tube simulations. Error bars are highlighted as black bars, within the symbol size **d.** and **e.** Drift velocity of the individual simulations trajectories for globular (light blue) and helical (light green) conformations. Average drift velocity is indicated by a red line along with a fitted curve (black line).

A more quantitative analysis of the over 1000 collision events observed in our simulations is shown in Fig. 3b. Here, each dot represents one of the collision events, separated according to the nearest approach (‘scattering parameter’), duration of the event, and velocity change (color). The three types of collisions form clusters, which are, however, not clearly separated from each other, and rather blend into each other, displaying a gradual range of these properties. Overall, the elastic collisions are short-lived, as are the swing-by events, but differ by both much shorter nearest approach and larger change of velocity (mainly direction). In contrast, due to the weaker intermolecular interaction, the velocity change of the swing-by events is smaller and scatters over a broader range. The adsorption/re-emission events are characterized by the closest approach, naturally, and by a broad spectrum of residence times ranging from 2 up to several 100 nanoseconds. Fig. 3c shows the total numbers of observed collisions for the globular and helical conformers; in line with its larger estimated CCS, significantly more collisions are seen for the helical conformations. An analysis of the root mean squared deviation (RMSD) of the simulated structures during and after the collisions (0.07±0.03 nm versus 0.12±0.04 nm, mean and standard deviation for the globular and helical conformations, respectively) shows that the impact of the gas molecules leaves the peptide structures largely unaffected and in particular does not trigger conformational transitions between helical and globular structures.

Next, we quantified the acceleration of the P1 peptide during the simulation by the applied (static) electric field against the increasing air resistance for the two conformations (Fig. 3d,e). Due to the relatively few collisions experienced within the highly diluted gas during each simulation, the individual traces (transparent lines) show considerable ‘Brownian motion’ scatter. Yet, the average velocity over 70 trajectories each (red solid lines) is well converged and follows the analytical solution of the Newtonian equation of motion for an accelerated object with air resistance proportional to its squared velocity (black lines, see Supplement); this analytical solution was fitted to the average velocity with the ratio between electric field strength and air resistance (*γ*) as the only fit parameter. A rapid initial velocity increase is seen, with a rate determined only by the peptide mass, charge and electric field strength (red dashed lines), and subsequently with decreasing acceleration towards a terminal drift velocity measured in the experiment. In line with the smaller number of collisions seen for the globular conformation (Fig. 3c), its terminal velocity (62.12±2.74 m/s), determined from the analytical fit at infinite time, is markedly larger than that of the the helical conformation (54.21±2.09 m/s).

For comparison, we estimated the drift velocities in the experiment from K0 values given the conditions used in the simulations (pressure: 2.7 mbar, temperature: 305 K and electric field: 20 V/cm) and obtained 78.31±1.75 m/s and 59.79±0.65 m/s for the two measured ion mobility peaks (peak value ± half width). Because particularly the pressure and density within the drift tube cannot be measured very accurately, this deviation is not unexpected, and one would therefore assume that the calculated drift velocity differs from the one estimated from the experiment by a common factor. Indeed correcting, e.g., the pressure to 3.2 mbar yields an estimate of 65.4 and 50.0 m/s, respectively, which lies very close to the values from MD. The main result here is that the extended conformation indeed shows a significantly slower drift velocity (and a correspondingly larger CCS) than the more compact conformation. In particular, the difference is large enough to explain the bimodal drift velocity distribution.

### Geometry-inspired approximation explains the bimodality at large scale

Our MD simulations suggest that globular and helical structures are stable in the drift tube environment, consistent with experimental results for poly-alanine peptides^15^. To further address the question if these two types of conformations are sufficient to explain the bimodal large-scale proteomics data, and since MD simulations for all peptides measured in the proteomics data are computationally quite demanding, we developed approximate treatments.

The, so called, geometric fit achieves a quantitative description of the large-scale proteomics data by statistically describing the dataset as a combination of two overlapping populations. The fit uses a joint model over all charge states where the CCS-mass distribution for each charge is parameterized as the sum of two independent densities, one for the globular and one for the helical population (see Online Methods and Fig. 4b). The helical population follows a linear CCS-mass relationship, as peptides grow primarily through helix extension. The globular population exhibits a power law dependence, CCS ~ mass^2/3^, reflecting uniform volume growth with added mass. Fig. 5b presents the geometric fit for charge state +3, while Supplementary Fig. 7 provides the fit results across all charge states, along with the associated error. This simple yet effective model closely aligns with the data, supporting the hypothesis that these two conformations indeed underlie and explain the observed bimodal distribution.

**Figure 4:**
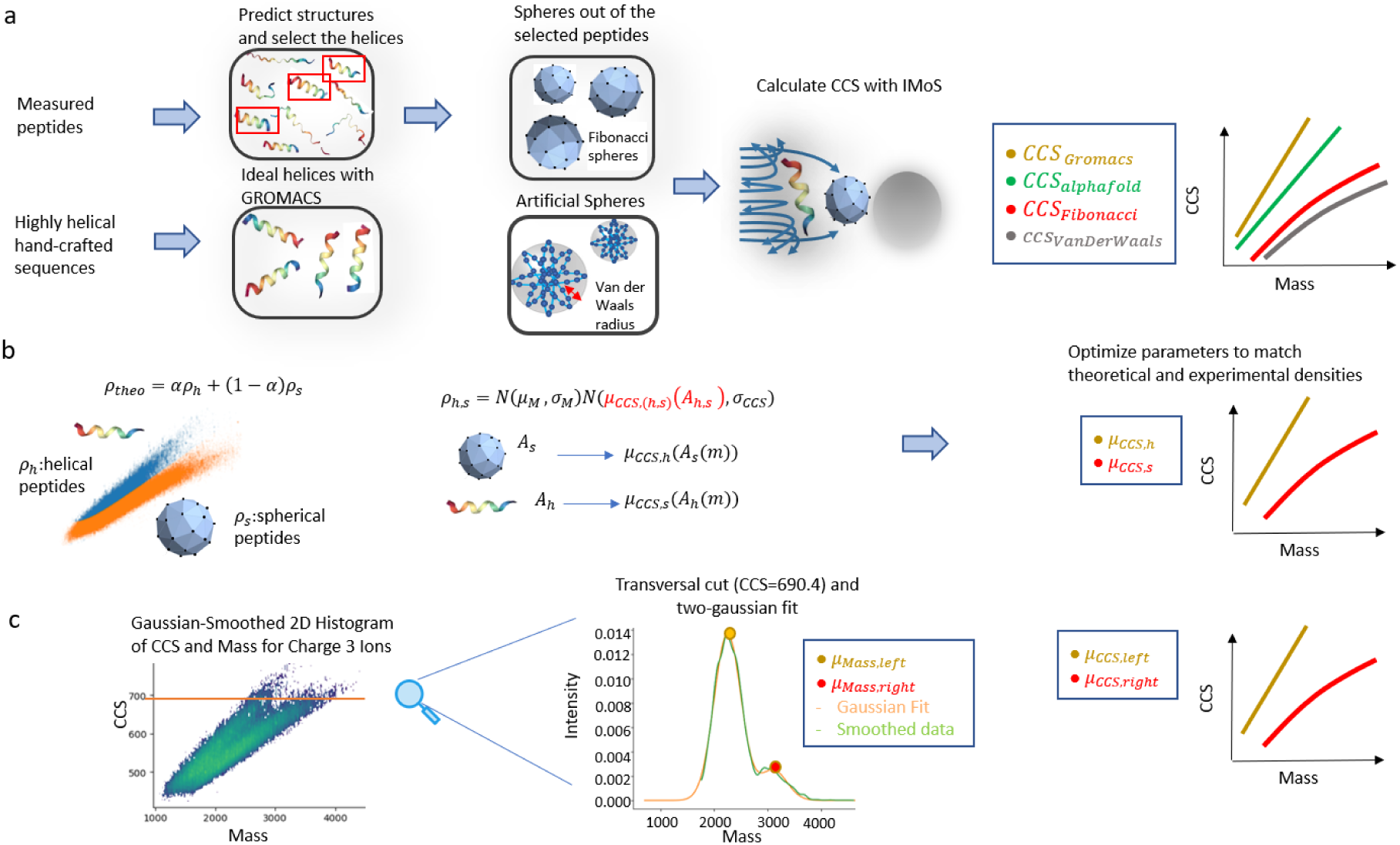
Fitting Models. **a. Geometric scattering**: Measured peptides undergo structure prediction with AlphaFold2, from which helical structures are selected. Spherical structures are generated from these helices using the Fibonacci sphere procedure. Hand-crafted sequences are converted into ideal helices with PyMOL and into spheres based on Van der Waals radii. CCS is then computed using IMoS for different structural models. **b. Geometric fit**: The large CCS vs mass dataset is modeled as a weighted sum of two Gaussian-distributed conformational states: helical peptides (*ρ_h_*) and spherical peptides (*ρ_s_*). The center of each Gaussian depends on the projected area of the peptide structure, which scales with mass. Parameters are optimized to best match theoretical and experimental densities. **c. Empirical fit:** A Gaussian-smoothed 2D histogram of experimental CCS vs. mass values is analyzed. A transversal cut at a fixed CCS (690.4 Å²) is shown, where a two-Gaussian fit distinguishes different conformational states. By repeating this procedure for all the CCS values while saving the fitted means per-population trend lines are generated.

**Figure 5:**
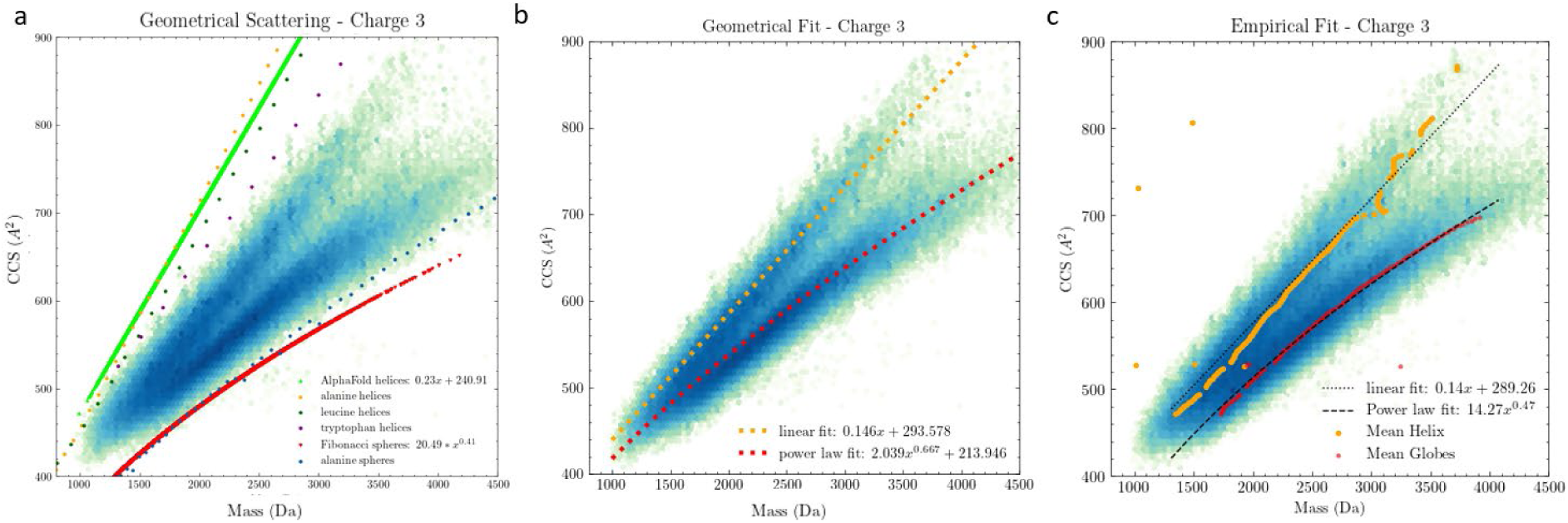
Comparison of CCS fitting approaches for charge state 3. **a. Geometric Fit:** A direct mathematical fit is applied to the experimental CCS vs. mass distribution. Both a linear fit (dotted red line) and a power-law **fit** (dashed orange line) are used to describe the trend in the data. **b. Geometric Scattering:** CCS predictions are derived from structural models, including AlphaFold2 helices, ideal helices, and spherical models (Fibonacci and Van der Waals spheres). These models define theoretical upper and lower CCS bounds (solid green and red lines), providing a structural basis for understanding the CCS-mass relationship. **c. Empiric Fit:** Experimentally derived trend lines are extracted by identifying the mean CCS values of helical peptides (orange) and globular peptides (red). A linear and a power-law fit are applied to characterize the observed experimental trends.

The intercept with the CCS axis (0.59 nm^2^) represents the charge-dependent contribution to the CCS corresponding to the value a massless unit charged peptide would exhibit. The helical slope (1.34 nm^2^kg^-1^mol^-1^) reflects the average CCS growth rate for a helical peptide as more amino acids are added. This procedure also provides the probability of a particular peptide belonging to either population. After calculating these probabilities, we found that for charge states +2, +3, and +4, the percentage of peptides in the upper (helical) population is approximately 4%, 20%, and 81%, respectively. Despite the good agreement, systematic errors remain as indicated by the non-random distribution of residuals (see Supplementary Fig. 5). Potential sources include deviations from sum of Gaussians of the data points and, particularly at lower masses, deviations of peptide conformations from the assumed globular or helical shapes.

Next, we determined whether Monte Carlo-based estimation of CCS values of ideal spheres and helices using IMoS^32^ can qualitatively describe the dataset (see Fig. 4a). Helical structures were obtained from AlphaFold2 predictions of experimentally measured peptides^33^ and compared to idealized polyalanine, polyleucine, and poly-tryptophan helices of varying lengths generated using PyMOL. Lacking well-defined globular candidates, we generated them via (i) transforming helices using the Fibonacci sphere method and (ii) distributing atoms on a spherical shell at van der Waals separations (see Online Methods). These structures were then evaluated in IMoS under frozen geometries, excluding partial charges. While this approach does not aim to be fully accurate, it successfully distinguishes the characteristic CCS trends of helical and globular structures. As shown in Fig. 5a, the simulated CCS values follow distinct scaling laws, with globular structures conforming to a power-law and helical structures with a linear trend. This clear separation supports our hypothesis and confirms that this simple geometric model qualitatively describes the observations. Quantitative deviations are seen for the exponent (2/3 from geometric considerations vs. 0.41 determined from data) as well as a systematic positive vertical offset, which both are likely due to the neglect of intermolecular interactions such as Coulomb and van der Waals forces; as seen in the atomistic drift tube simulations, these interactions affect the scattering processes and thus also the CCS.

Finally, we performed a solely data-driven fit, here referred to as empirical fit, designed to optimally separate the two populations for use in CCS prediction. In this approach, we divided the (CCS, Mass) space into bins, smoothed the data using a Gaussian kernel, and assumed that each slice at a constant CCS could be modeled as a mixture of two Gaussians (see Online Methods and Fig. 4c). Selected slices for charge state +4 are shown in Supplementary Fig. 8. Fig. 5c shows the mean of both Gaussians for all the transversal slices of the charge state +3 dataset (yellow dots for the left Gaussian, red dots for the right Gaussian), overlaid on the measured data. The means accurately trace the regions of highest density for each population and closely follow linear and power-law trends, consistent with the expected structural scaling behaviors. This purely data-driven method yields probability distributions per population and per slice without relying on prior assumptions about peptide geometry. Importantly, it allows us to assign labels to peptides based on the inferred probability distributions using their CCS, charge and mass across the whole dataset. These labels allow us to train per-population regressors for CCS prediction.

### Two-valued machine learning prediction improves identification of peptides in proteomics

Existing mobility predictors are typically trained to predict the mobility of a peptide’s most intense feature, overlooking the fact that many peptides exhibit two distinct, well-defined mobility values due to the existence of two stable conformations. This oversight introduces stochasticity and may reduce peptide identification rates by predicting the ‘wrong’ mode for a peptide in a given dataset. To address this issue, we divided the training set into two clusters by assigning each peptide a probability of belonging to either cluster using the empirical and geometric fit described in the previous section and selecting the most likely one (see Methods). We then trained a bidirectional recurrent neural network (RNN) on the encoded sequence and charge for each cluster across all charge states. Similarly, we derived sequence-based features, combined them with metadata, and trained a XGBoost regressor for each stable conformation. As a baseline, we trained both models on the unseparated dataset, representing the conventional approach used in prior studies. To benchmark the performance of these models, we employed two approaches: evaluating prediction error on an independent test set and assessing peptide identification rates in a data-independent acquisition (DIA) experiment using predicted libraries.

We tested the models on an independent dataset from the ProteomeTools project and measured the relative error with respect to the most intense value within each cluster. The results for charge 3, shown in Fig. 6a, reveal that the RNN with empirical labeling outperforms other models in the most bi-modal case, as expected from its optimal separation of peptide populations. The RNN with geometrical-fitting labeling ranked a close second, showing that it also successfully learned the bimodality of the data. The baseline RNN trained on the unseparatrd dataset performs similarly to the XGBoost models trained with labeled data in a basic feature space derived from amino acid counts, clearly exhibiting the importance of the proper labeling. Notably, the poor performance of the XGBoost model for charge state +3 in the unlabeled case highlights the limited flexibility of the feature space, as this charge state is particularly challenging to predict.

**Figure 6:**
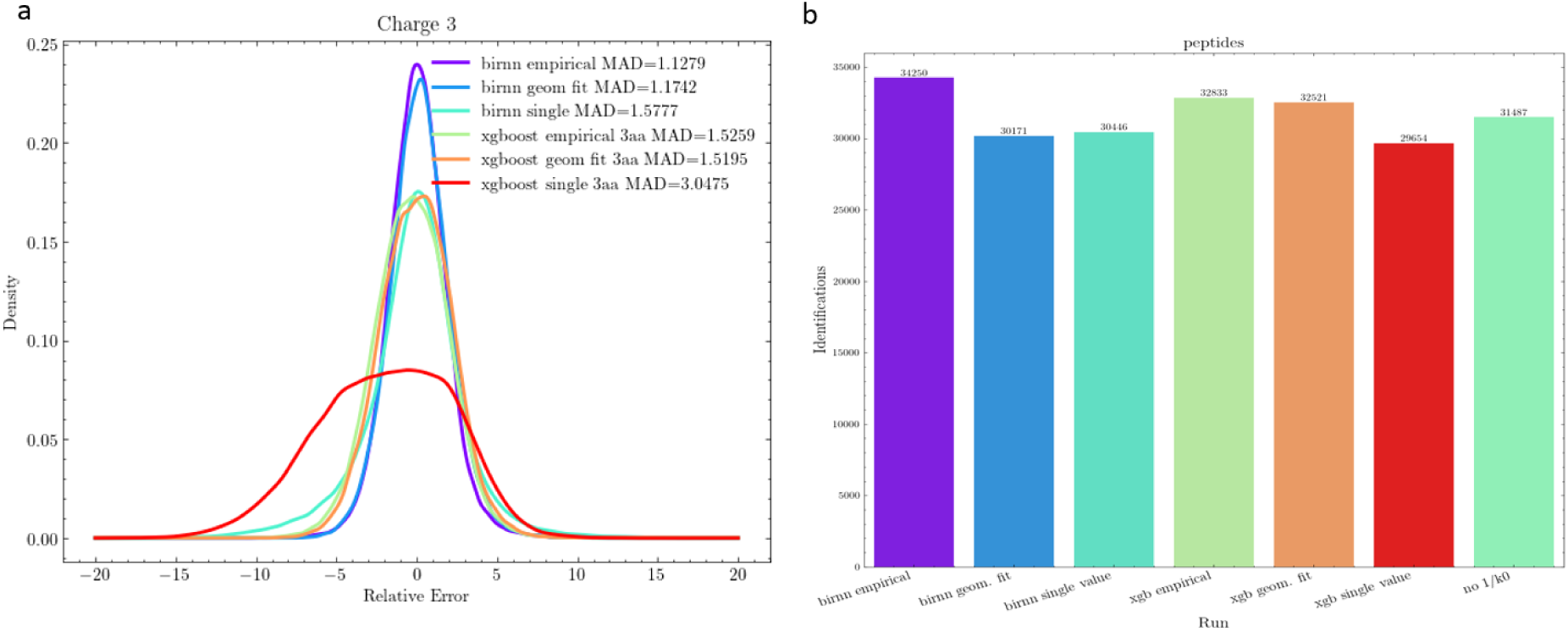
Benchmark of mobility predictors. **a. Relative Error in Test Set:** Comparison of two machine learning models (Bi-RNN and XGBoost) using three different peptide-labeling methods (empirical fitting, geometrical fitting, and baseline with no labeling). The relative error of their predictions is evaluated on the ProteomeTools dataset. **b. Peptide identifications:** The same models and labeling methods are assessed based on the number of peptide identifications in a spectral library used for a DIA run with MaxDIA.

Lastly, we used spectral libraries predicted with DeepMass^34^, predicted the reduced mobility of the existing sequences and tested them in a data-independent acquisition (DIA) experiment. Fig. 6b shows the number of identified peptides using libraries generated by different models. The RNN with data separated by the empirical fit clearly outperformed all other approaches, followed by XGBoost with empirical fit labeling and XGBoost with geometric fit labeling. This result underscores that for mobility prediction, the separation of peptide populations is more critical than the choice of model itself. Interestingly, omitting mobility values ranks fourth, potentially because MaxQuant bypasses mobility filtering in such cases. However, this result also demonstrates that poor mobility predictors can severely impact performance. The RNN with geometrical fit labeling and the baseline RNN without labeling showed similar performance, followed by the baseline XGBoost model as the least effective.

## CONCLUSIONS

Our combined approach involving ion mobility spectrometry measurements, atomistic molecular dynamics simulations, and geometric modelling, revealed that the bimodality in peptide CCS data as it is produced in proteomics data arises mainly from conformational heterogeneity of a larger peptide population, and specifically from the presence of globular and helical conformations. Our molecular dynamics simulations of selected peptides in gas phase show a strong prevalence for these two configuration types for peptides. Full simulations of peptides mimicking the drift tube experiments show that the differences in drift velocities between compact and extended conformations suffice to explain the observed bimodal drift velocity distribution. We found not only elastic collision events, as one might have expected, but also adsorption/re-emission and swing-by events, which, combined, contribute to the observed CSS. Accordingly, our simulations can serve to refine CSS estimates. Based on our simulations, we have successfully derived more approximate yet efficient models which can be successfully applied to the large-scale data. Explicit inclusion of the bi-modality in machine learning-based CCS prediction turned out to markedly improve prediction accuracy.

## ONLINE METHODS

### Data download and processing

We downloaded and reprocessed the dataset described in Meier et al.^35^ using MaxQuant^36^ v2.6.6.0. The search engine Andromeda^37^ was used for peptide identification by matching the measured spectra to theoretical spectra generated via in-silico digestion of reference proteomes with specific enzymes (trypsin, LysC, or LysN). Cysteine carbamidomethylation was set as a fixed modification, while oxidation of methionine and protein N-terminal acetylation were set as variable modifications. Additionally, a list of 245 potential contaminants was included in the search. The FASTA files of the reference proteomes, including isoforms, were downloaded from UniProt (release 09/2023) and contained the following: Homo sapiens (103,830 proteins), Saccharomyces cerevisiae (6,091 proteins), Drosophila melanogaster (23,543 proteins), Escherichia coli (4,415 proteins), and Caenorhabditis elegans (28,540 proteins). Following the original publication, for the five-species dataset, the maximum mass tolerances were set to 20 ppm for precursors and 40 ppm for fragment ions. Each set of synthetic peptides was analyzed in independent MaxQuant runs, with libraries generated in silico by tryptic digestion of the human proteome. The additional HeLa dataset was processed as outlined in Meier et al. (PMID:30385480). The ‘TIMS half width’ was set to 4, and the ‘TIMS mass resolution’ to 32,000. The maximum mass tolerance for precursors was set to 70 ppm, for fragments to 35 ppm, and for precursors after recalibration to 20 ppm. The diaPASEF dataset was analyzed, as well, with MaxQuant v.2.6.6.0 using spectral libraries predicted by DeepMass^34^. The mobility values were predicted with the different regression models and added to the libraries manually.

### Analysis of the MaxQuant output

The MaxQuant output was analyzed using Python 3.7.11 with the NumPy, Pandas, SciPy, and Matplotlib libraries. We filtered out decoy peptides, potential contaminants, features with null intensity, and peptides identified with only one positive charge. To integrate all the different MaxQuant runs into a single dataset, systematic offsets were corrected through a machine-learning-based approach. Specifically, we designated the larger HeLa dataset as the master run, trained a recursive neural network on its most intense feature per precursor, and predicted the mobility values for the most intense features of precursors in the other experiments. The difference between the predicted and measured values was then calculated, grouped by raw file, and summarized as a median correction factor for each file. This approach remained robust even when overlap with the master run was limited or nonexistent, such as when LysN was used as the protease.

### Geometric fit

For the geometrical fit, we constructed 2D histograms in the (CCS, Mass) space for each charge state, *ρ_m_*. We modeled the total density as the sum of two bi-dimensional Gaussian distributions with relative abundances, *ρ_th_* = *αρ_h_* + (1 − *α*)*ρ_s_*, where *α* is the abundance of the helical population and *ρ_h_*, *ρ_c_* are gaussian distributions representing the helical and spherical populations, 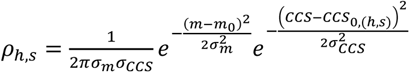. The center of each Gaussian in the CCS dimension was assumed to follow a linear relationship with the projected area, which itself is determined by volume and, consequently, mass, where 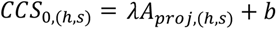. For the lower population, we assumed a spherical geometry for the projected area, 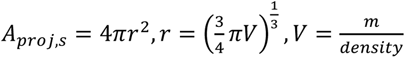, while for the upper population, a cylindrical 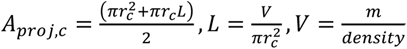. The density parameter, initialized to 1000 kg/(mol nm^3^), was optimized during the fitting process through a scaling factor. Gaussian widths were optimized but shared across conformations, as were the parameters for the linear model of the center in the CCS dimension. The relative abundance of each Gaussian was modeled as a linear function, with parameters optimized across populations, *α* = *γm* + *β*. Additionally, the cylinder radius *r_c_* was treated as a free parameter. Once fitting was complete, we computed the probability of each point belonging to the spherical population and labeled it as spherical when this probability was ≥ 0.5.

### Empirical fit

For the empirical fitting, we calculated the conditional probability density P(CCS|Mass) for each charge state and performed transversal slicing at constant CCS values. Each slice was smoothed using a Gaussian kernel with sigma=1 Da for net charge two and sigma=4 Da for net charges three and four. Then we fitted a model consisting of the sum of two Gaussians to the resulting distribution. For charge two, Gaussian heights were constrained to (0, 1.0), means to (900, 3200), widths to values greater than 50. For charge +3, the widths were allowed to differ but retained the same minimum threshold, the means were constrained to (1000, 4000) and only slices with CCS<700 were considered as the low point density above this level made the fitting unstable. For charge +4, the mean range was restricted to (1500, 4500), the maximum CCS for the left population was 900 and for the right one 800. Initial parameters for the fitting process were determined using a peak-finding algorithm with thresholds set to a minimum height of 0.0005 and minimum prominence of 0.0001. The two most prominent peaks were used as starting points; if only one peak was detected, it was duplicated, and slices without peaks were skipped. To reduce the noise, an ad-hoc condition was applied, that the fitted means had to differ at least by 200 units. The fitted parameters were sorted based on their means to construct probability density functions for each population within each CCS bin. The left and right Gaussian means were extracted per slice and fitted using a linear model for the left population and a power-law model for the right. Based on these probability density functions; points were assigned to the population with the highest probability.

### Geometric scattering

For the geometric scattering we generated ideal spherical and helical peptides. To generate idealized spherical peptide models we used two methods, the Fibonacci sphere method and our own algorithm based on placing atoms on a spherical shell separated by the Van der Waals radius. First we utilized Fibonacci spheres, a mathematical approach for evenly distributing N points on the surface of a sphere. This method allowed for the sequential placement of protein atoms on the spherical surface, ensuring uniform distribution. To construct non-hollow spherical models, we generated 4 concentric Fibonacci spheres with progressively smaller radii. The radii were defined based on the framework provided by^38^, which describes the minimal radius of a spherical protein that contains a given mass. This approach enabled the systematic construction of densely packed ideal spherical protein structures. We calculated the spherical conformations for a total of 60496 peptides, comprising 44093 peptides with a charge of +2, 14665 peptides with a charge of +3, and 1738 peptides with a charge of +4. We adapted the radii to smaller values as has been used in the reference^38^, 80% of it for charge +3, +4 and 70% for charge +2 which can be attributed to the distinct properties of the gaseous environment, where the absence of solvent effects and amplified electrostatic repulsion due to the lack of dielectric screening influence the conformation. In the second method, we considered spheres with radii ranging from 0.5 to 1.1 nm and assumed a fixed density of 1000 kg/m³. We then determined a dense spherical grid with equal angular steps where the largest separation (at the equatorial) is determined by half of the hydrogen van der Waals radius. Finally, we added atoms to the grid points from a list of atom names representing the composition of alanine if it is not overlapping with already placed atoms. The resulting atom names and coordinates were used to construct a PDB file. To generate the ideal helical peptides, we employed AlphaFold2^33^ and PyMOL (Version 3.0, Schrödinger, LLC). When using AlphaFold2 we predicted the solution structure of the measured peptides. Subsequently, we analyzed the secondary structures of these peptides and selected those that exhibited a consistent alpha-helical structure along the peptide backbone. We detected helical conformations for a total of 60496 peptides, comprising 44093 peptides with a charge of +2, 14665 peptides with a charge of +3, and 1738 peptides with a charge of +4. For comparison, we also included helical structures with ideal dihedral angles generated using PyMOL for sequences ranging from 7 to 40 amino acids with high helical propensity, such as alanine-, leucine-, and tryptophan-based peptides. The CCS values of the peptides were calculated using the Ion Mobility Spectrometry Suite (IMoS) software, version 1.10. Default parameters were employed, with the exception of the pressure, which was set to 270 mbar, and the temperature, which was adjusted to 305 K, and the respective charges to replicate the experimental conditions. The CCS values were determined using the Trajectory Method Lennard-Jones (TMLJ) approach, which calculates the momentum exchange between the buffer gas and the peptide by simulating individual trajectories of gas molecules. This method also accounts for interaction potentials between the buffer gas and the molecule. The TMLJ method employs a 4-6-12 potential with optimized Lennard-Jones parameters, making it the gold standard for accuracy in CCS calculations (source IMoS).

### MD quenching simulations

We performed fully atomistic molecular dynamics temperature quenching simulations *in vacuo* to sample and predict the conformation of the measured peptides. As unbiased starting structures, fully extended conformations were generated using Pymol (Version 3.0 Schrödinger, LLC.). Although the total charges of the peptides are known, in many cases their distributions are ambiguous. To avoid this uncertainty, sequences were selected where the number of arginine, histidine, and lysine residues together with the charged N-terminal sums up to the expected total charge, such that this charge distribution is uniquely specified. In two other cases, the two most plausible different charge states (among the many other combinatorial possibilities) were used as shown in Table 1. Aspartate and glutamate residues we usually protonated and therefore, neutral. The extended peptides were placed in a cubic box with a side length of 100 nm. Energy minimization (steep integrator, 3000 steps) and a four step equilibration was performed with increasing time steps: 0.1 fs, 0.5 fs and 1 fs in NVT and lastly in an NPT ensemble. The resulting structures were used for the quenching simulations where a simulated annealing temperature quench was applied to allow the peptides finding their equilibrium conformation. Specifically, the initial temperature was 600 K and maintained for 10 ns, then linearly and slowly decreased to 305 K over a time period of 500 ns and subsequently kept at 305 K for another 40 ns to facilitate thermal equilibration. Due to the absence of solvent and the presence of a net charge, we changed the simulation parameters relative to those typically used for conventional simulations as follows. i) Double precision compilation of GROMACS (version 2023.4 10.1016/j.softx.2015.06.001) was necessary and a 1 fs integration time step; ii) cutoff distances were set to 30 nm, such that all peptide atoms interacted with all other atoms explicitly via Coulomb and Lenard-Jones forces; accordingly, a long neighbor search interval (1 ns) was used, and (iii) no Particle Mesh Ewald (PME) method was required, which would fail due to the total net charge of the system; iv) no pressure coupling was applied. The system was periodic, but the center of mass was kept at the center of the simulation box. The temperature was controlled by the V-rescale algorithm^39^. All simulations were performed with the CHARMM36m force field^40^ with added oxygen and nitrogen molecule parameters adapted from Wang et al.^41^ for the later drift tube simulations. The quenching procedure was repeated 1000 times for every peptide with starting velocities chosen randomly and different in each case from a Boltzmann distribution. All atomic positions were saved every 10 ns.

### MD drift tube simulations

From the quenching simulations of peptide P1, 7 helical and 7 globular conformations were selected as starting structures to simulate the full drift tube environment. As in the experiment, an air mixture was used as the inert gas at 2.7 mbar (the approximate pressure estimated in the experiments) and 305 K, which translated into 51 N_2_ and 13 O_2_ molecules in the cubic simulation box of 100 nm side length. This ‘drift tube’ simulations box was initially created and simulated without the peptide for 100 ns at 305 K to obtain an equilibrated system, to which subsequently the peptide structures were inserted. Starting velocities were taken from the appropriate previous simulations. In each drift tube simulation, an electric field of 20 V/cm parallel to the x-axis was applied.

The aim of this simulation was to test the effect of conformation on the velocity increase and the terminal velocity of the peptide that results from the balance between the force exerted by the electric field and the collisions with the gas molecules. Therefore, center of mass motion removal was applied only to the air molecules in the box (once every ns), and not to the moving peptide. To account for the need to maintain the temperature of the air while not interfering with the velocities of the peptide atoms, we used V-rescale temperature coupling separately for the gas (with a coupling constant *τ* = 5 ps) and for the peptide (*τ* = 1 ps). The extremely long coupling time for the peptide ensured a neglible effect of the heat bath. Temperature coupling of only a subset of the simulated atoms is currently not possible with GROMACS. As for the quenching simulations, large cutoffs were used without PME, but due to the fast-moving gas molecules the neighbor search was performed for each integration step (1 fs). Atomic coordinates and velocities were recorded every nanosecond. Each of the 14 peptide conformations was simulated 10 times independently, each starting with a different set of random positions and velocities of the air molecules. In order to obtain the equilibrium drift velocity (*v*_d_) from these 10 MD simulations for each peptide, the analytical solution of the Newtonian equation of motion for an accelerated object with air resistance proportional to its squared velocity

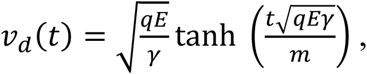

was fitted to the time-dependent average velocity obtained from the simulations, with the ratio between electric field strength and air resistance (*γ*) as the only fit parameter. Here, *t* is time, *q* is the peptide charge (+3), *E* is the electric field strength (Vm), *m* is the mass of the peptide (kg/mol), and *γ* is defined as

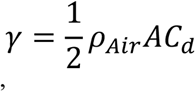

with gas density *ρ*_Air_, surface area *A* of the peptide, and shape parameter *C*_d_. The air resistance *γ* includes both shape and surface area differences between peptide conformations, lumped up into one single fit parameter.

### Electronic structure analysis with Gaussian

We analyzed the energy of a peptide for multiple runs obtained from molecular simulations, which display both globular and helical conformations, to determine whether there is an energetic disparity between these secondary structures. Using Gaussian software, we performed geometry optimizations by applying density functional theory (DFT) with a ωB97X-D functional and the 6-31G(d) basis set. To reduce computational costs, we used the NoSymm keyword to prevent molecular reorientation and geom=connectivity to explicitly define the molecular connectivity, a commonly used approach for such optimizations. Additionally, we applied the SCF=NoVarAcc option to accelerate the convergence behavior of the self-consistent field (SCF) calculations. The optimizations did not converge to the default convergence criteria of Delta E = 10^-6^ atomic units but sufficient to enable comparison between the two conformations. During the optimization process, the secondary structure for the different runs was maintained.

### Machine learning predictions

For each considered separation, we trained a bi-directional recurrent neural network (RNN) and an XGBoost^42^ regression model. As a baseline, we also trained both models on the most intense feature per precursor without applying separation. For the deep learning model, peptide sequences were encoded to include modifications, resulting in 26 unique classes. The encoded sequences were padded to form matrices with 66 columns, and the net charge was appended as an additional column. The dataset was split into training (90%) and validation (10%) subsets, with reproducibility ensured by setting a fixed random seed (42). The encoded training sequences were passed through an embedding layer connected to two bi-directional LSTM layers, each layer with 128 units and a dropout probability of 0.5. Global pooling was applied along the sequence dimension after the final LSTM layer to ensure consistent shape across instances. The charge value was concatenated to the hidden state and fed into a fully connected layer with 128 neurons and a dropout probability of 0.4. ReLU activation was applied before the final output. The model was trained for 200 epochs with a batch size of 64 using an inverse square root learning rate scheduler (normalization factor = 1056) with a warmup phase of 10,000 steps. Optimization was performed using the Adam optimizer (β₁ = 0.9, β₂ = 0.98, ε = 1e-9). The final model, implemented in Python with TensorFlow (arXiv:1603.04467), contained 694,658 parameters and was used for both the separation and baseline cases. All computations were performed on an NVIDIA RTX 5000 GPU.

For the tree-based model, peptide sequences were encoded by concatenating sequence-derived features with metadata. Sequence features included amino acid counts, dipeptide, and tripeptide compositions. Metadata comprised peptide mass, mass-to-charge ratio, length, charge, and one-hot-encoded enzyme labels. This process resulted in feature vectors of length 18,285. The dataset was split into training (90%) and validation (10%) subsets with a fixed random seed (42) for reproducibility. Hyperparameters for the separation-specific and baseline cases were optimized using Hyperopt (10.1088/1749-4699/8/1/014008). The Python wrapper of the XGBoost library was employed for model training and predictions.

## Data availability

MaxQuant results and supplementary movies have been deposited at Mendeley Data under https://data.mendeley.com/preview/szrn5srhyw?a=3381e6af-4c79-4f92-be3a-9ca13ef0c9fa.

## Code availability

Custom code used for the data analysis has been deposited at https://github.com/cox-labs/CCS.

## ACKNOWLEDGEMENTS

This work was supported by the German Ministry for Science and Education funding action CLINSPECT-M [FKZ 161L0214E].

## AUTHOR CONTRIBUTIONS

J.R. did machine learning-based data analysis. D.S. performed the MD simulations, T.K. did the IMoS analysis, J.R., D.S. and T.K. did all other data analysis, J.R., D.S., C.W., H.G. and J.C wrote the manuscript, C.W., H.G. and J.C. directed the project. All authors read and approved the final manuscript.

