## Supplementary Figures for "Bimodal peptide collision cross section distribution reflects two stable conformations in the gas phase"

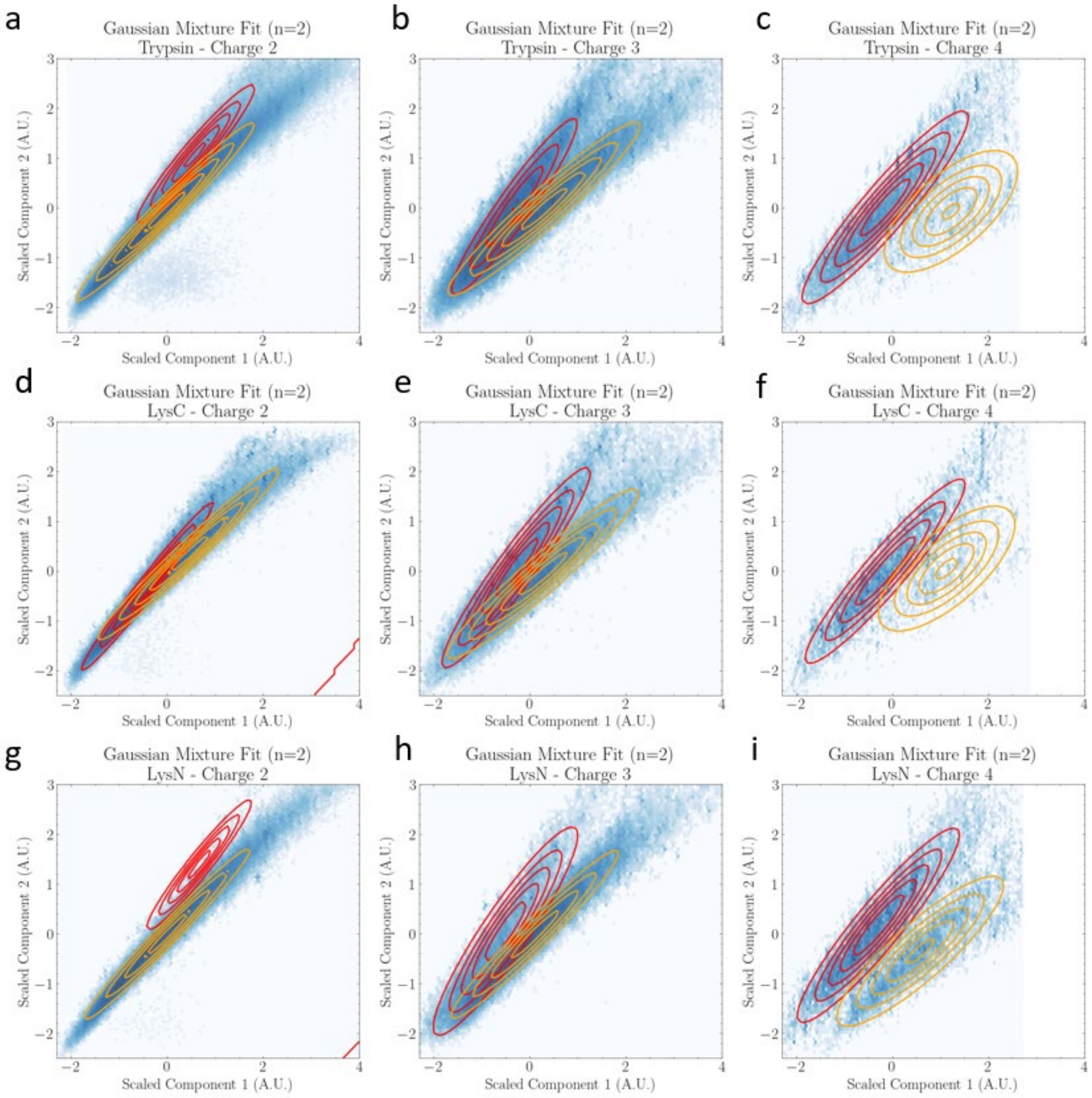


**Supplementary Figure 1.** Fit to the sum of two bivariate normal distributions per protease and charge state overlaid on the corresponding distribution. The distributions are normalized to have zero mean and unit variance.

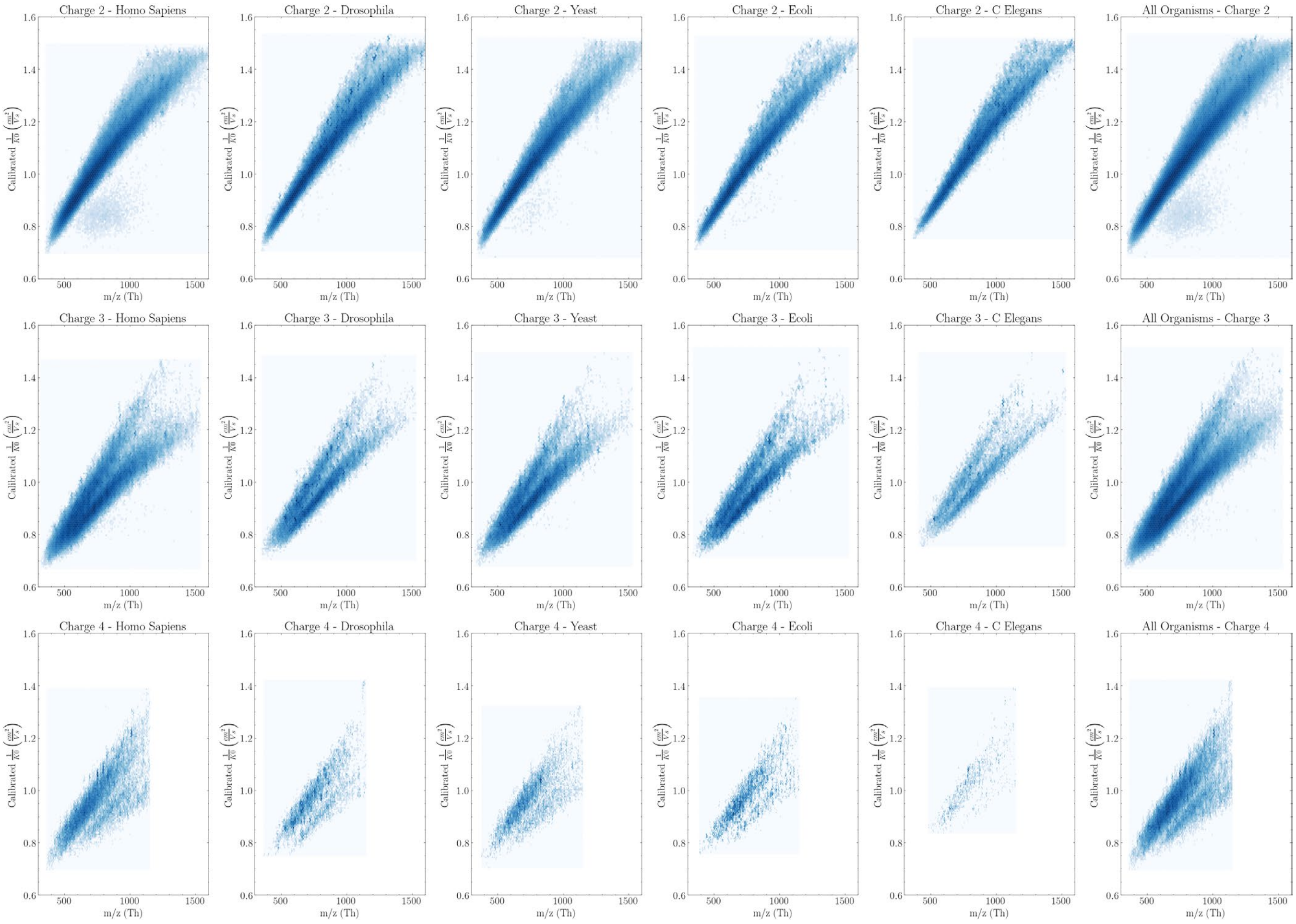


**Supplementary Figure 2.** Distribution of peptides in the space of reduced mobility calibrated across all runs versus mass-to-charge ratio for all charges and organisms.

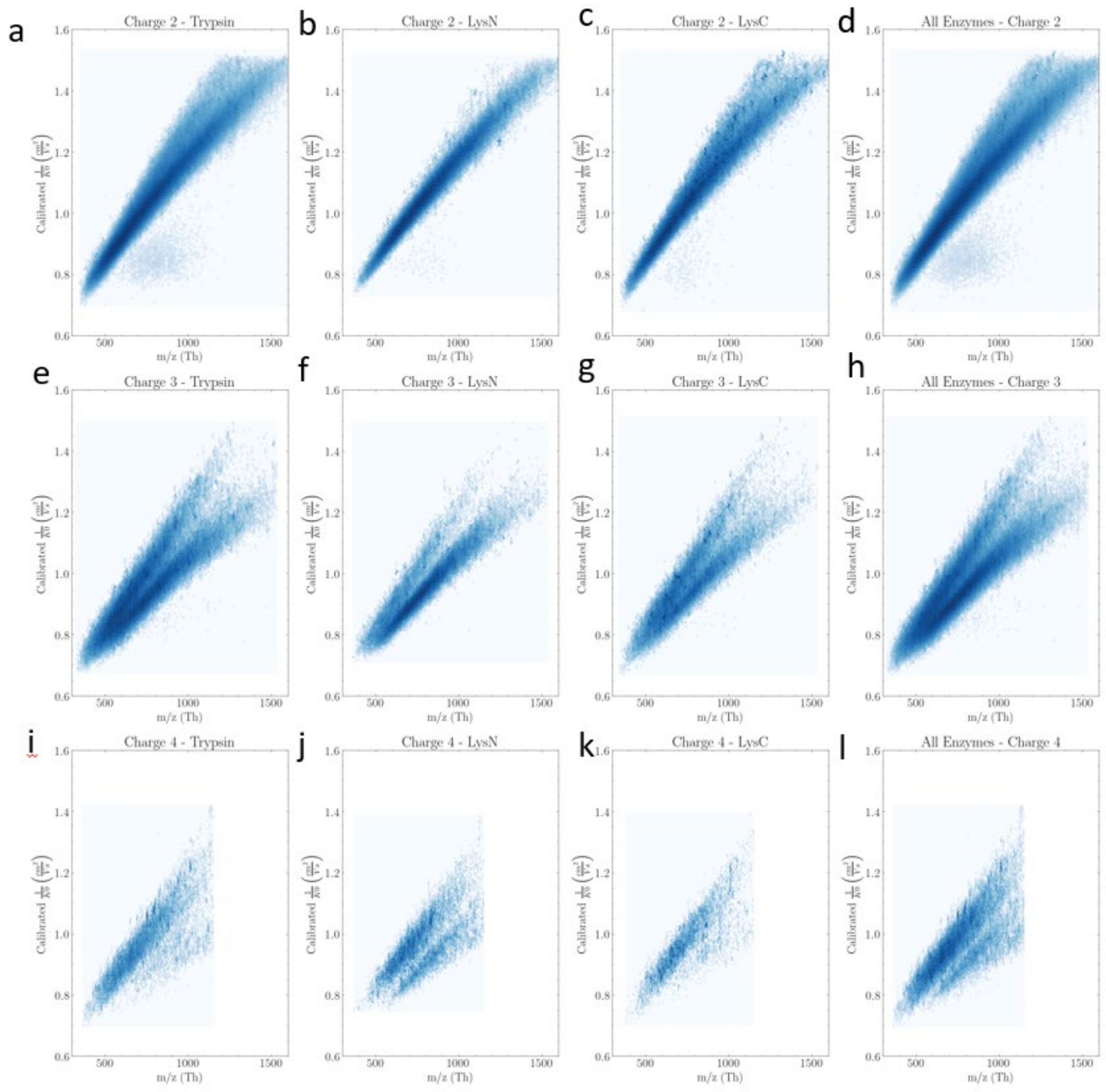


**Supplementary Figure 3** Distribution of peptides in the space of reduced mobility calibrated across all runs versus mass-to-charge ratio for all charges and proteases.

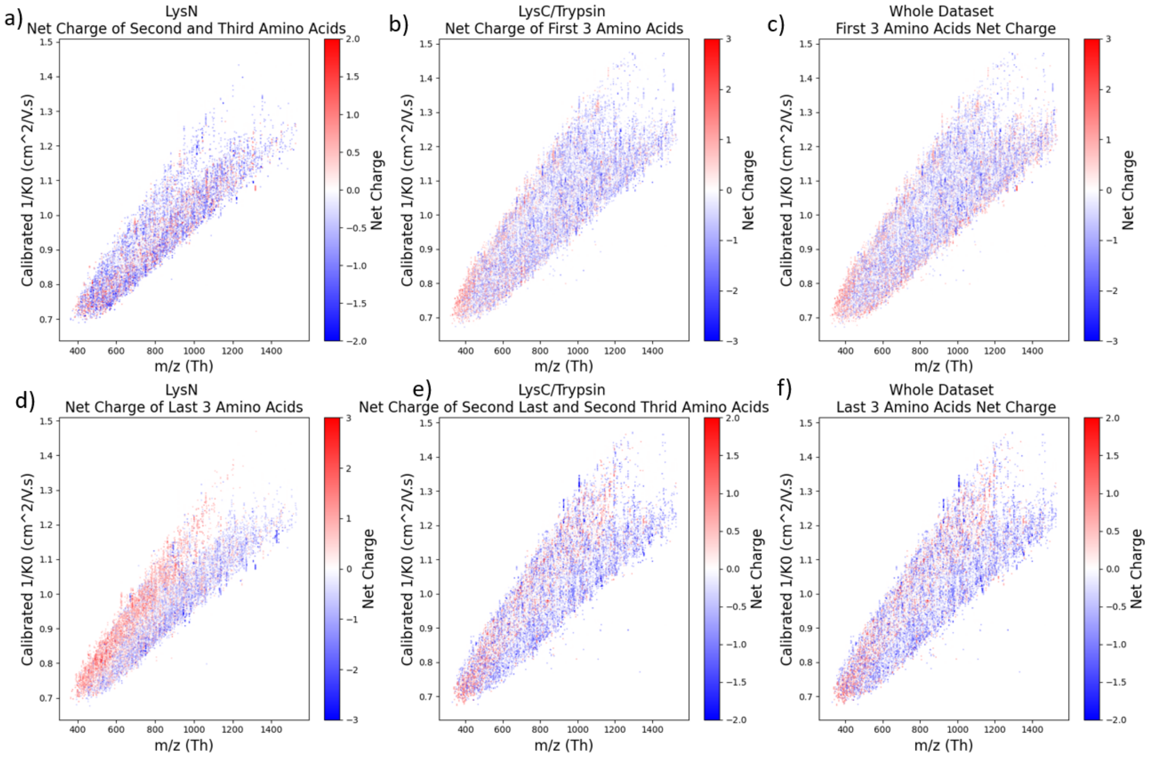


**Supplementary Figure 4.** Distribution of peptides in the space of reduced mobility calibrated across all runs versus mass-to-charge ratio for two enzymatic groups: LysC/Trypsin and LysN. Each distribution is colored by net charge in either the first three or the last three amino acids ignoring the charge associated to the protease. The distribution of the total dataset without enzymatic division is also shown on the right-most column.

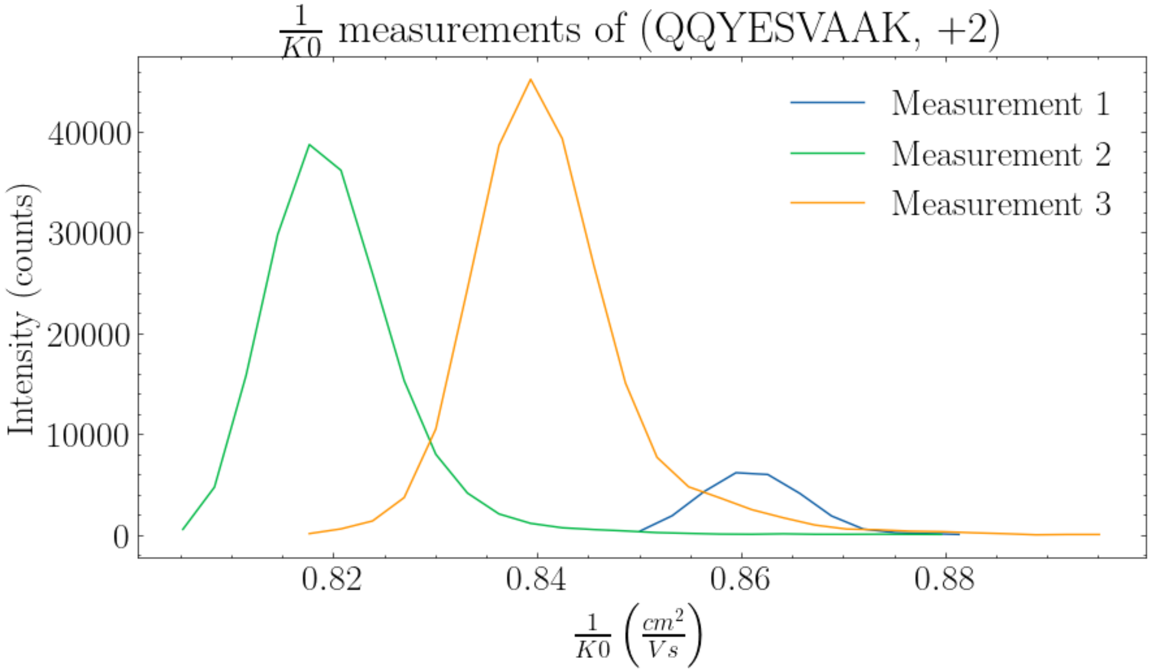


**Supplementary Figure 5.** Intensity profile on the reduced mobility dimension for a particular precursor (see title) that exhibits multiple peaks.

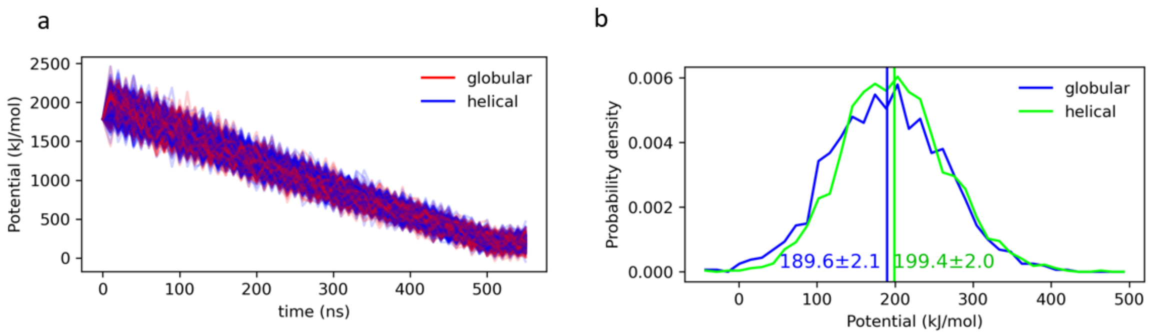


**Supplementary Figure 6.** Potential energy histogram of globular and helical conformations of the P1 peptide. Vertical lines and label indicate the mean potential energy and its standard error. The last 40 ns of the quenching simulations at constant temperature 305 K were used for this analysis. Grouping into globular (blue) and helical (green) was based on predicted CCS for the final conformation, structures above 850 Å2 were considered helical, otherwise globular.

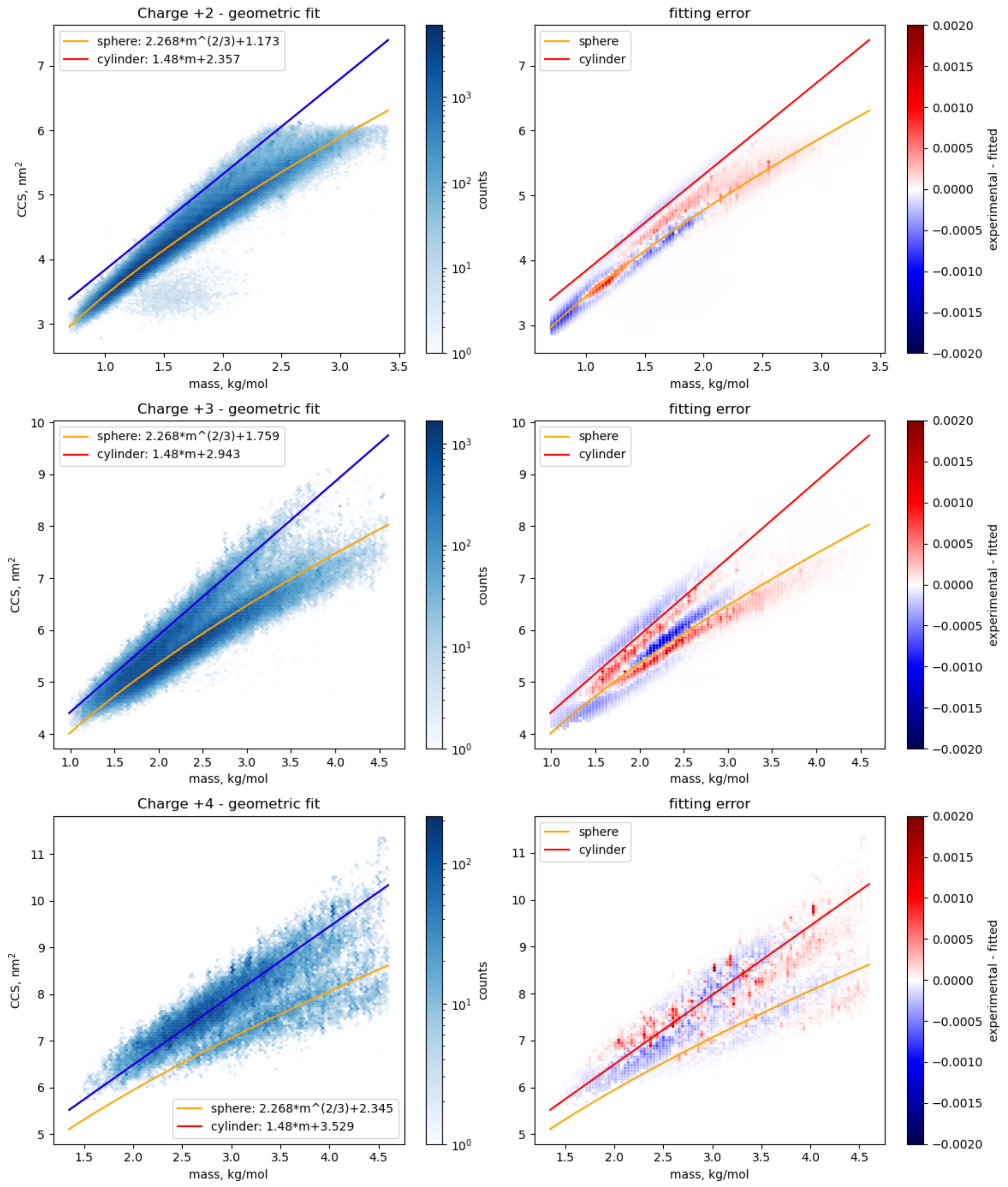


**Supplementary Figure 7.** Geometric fit (left column) and fitting error distribution (right column) for each charge state.

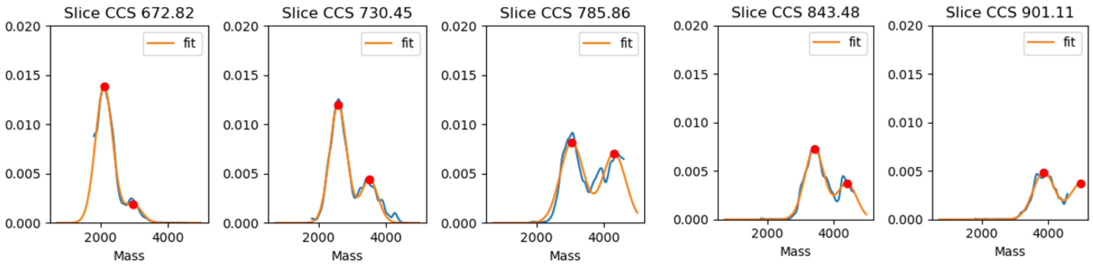


**Supplementary Figure 8.** Selected transversal slices with constant CCS over the mass dimension(blue line) for charge state four together with the fit to the sum of two one-dimensional gaussian distributions (orange line).

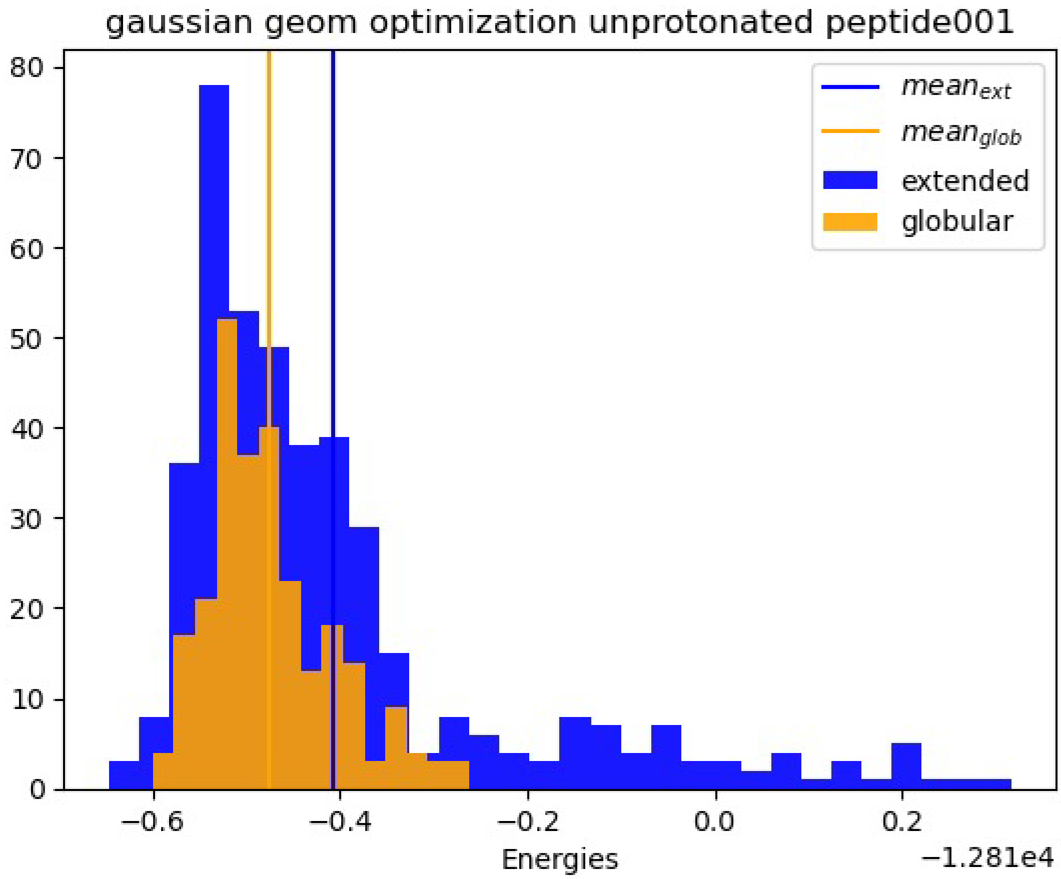


**Supplementary Figure 9.** Distribution of energies of peptide001(see table 1) in different configurations obtained by simulations. The extended or globular label was set by observation of the CCS distribution of the simulated peptides. The energies were calculated using the Gaussian software.
